# Antifungal activities of artesunate, chloramphenicol, and cotrimoxazole against *Basidiobolus* species, the causal agents of gastrointestinal basidiobolomycosis

**DOI:** 10.1101/2021.01.11.426312

**Authors:** Saleh Al-Qahtani, Martin R.P. Joseph, Ahmed M. Al Hakami, Ali A. Asseri, Anjali Mathew, Ali Al Bshabshe, Suliman Alhumayed, Mohamed E. Hamid

**Affiliations:** Department of Medicine, College of Medicine, King Khalid University, Abha, Kingdom of Saudi Arabia; Department of Microbiology, College of Medicine, King Khalid University, Abha, Kingdom of Saudi Arabia; Department of Child Health, College of Medicine, King Khalid University, Abha, Kingdom of Saudi Arabia; Community Health Centre, Vandanmedu, 685551, Kerala, India

**Author notes:** The authors contributed equally to this paper (Saleh Al-Qahtani, Martin R.P. Joseph).

**Keywords:** Trimethoprim/ sulfamethoxazole (Bactrim, Septrin), antifungal screening, Broth Microdilution, *Basidiobolus*, synergism, human gastrointestinal basidiobolomycosis

## Abstract

*Basidiobolus* species (n =13) isolated from human gastrointestinal basidiobolomycosis and lizards were tested against artesunate, chloramphenicol, and cotrimoxazole. The three agents exhibited inhibitory actions against *Basidiobolus* species comparable to the known antifungals. The combined effects of artesunate + voriconazole and cotrimoxazole + voriconazole have significant synergic effects, p = 0.003 and p = 0.021, respectively. These are promising results that enhance accelerated combined treatment of GIB in humans particularly the combination of artesunate and voriconazole.

## Introductions

Basidiobolomycosis, which is caused by pathogenic *Basidiobolus* species, notably *Basidiobolus ranarum*, is a rare fungal infection affecting the skin and gastrointestinal tract. Thhe disease is mainly reported from tropical and subtropical regions. *Basidiobolus* (Order: *Entomophthorales*) is a zygomycete filamentous fungus isolated from plant debris, soil, amphibians, reptiles, and insectivorous bats, and lice (1, 2). One of the severe forms of basidiobolomycosis is human gastrointestinal basidiobolomycosis (GIB). Most of the GIB cases were pediatrics and predominately reported from the south of Saudi Arabia (3–6).

Diagnosis of GIB based on clinical suspicion is challenging and requires histopathological and mycological confirmation. Similarly, treatment is difficult given the nature of the disease, which requires endoscopic examination, early surgical resection of the infected tissue, and prolonged treatment with antifungals. Antifungals such as itraconazole provide suitable treatment options (5). Many of the azole antifungals, for instance, itraconazole, can produce a range of serious cardiac and fluid-associated undesirable episodes. Dose decrease or cessation usually resulted in symptomatic improvement or reversal (7).

There is a continuous need for new antifungals because of the limited number of accessible therapeutic drugs for treating fungal infections. As a result of the widespread antibiotic resistance and lack of novel antibiotic options, studies are needed to discover the possible activities of known antibiotics to act on unconventional pathogens (8). Several studies have demonstrated a good clinical outcome of the antifungal effects of “non-antifungal drugs” against true pathogenic species (9–11), for example, Trimethoprim/ sulfamethoxazole (8, 9), artesunate (12), and chloramphenicol (11, 13). Cotrimoxazole is one of these “non-antifungal drugs,” which is widely used to treat a variety of bacterial infections. Trimethoprim/ sulfamethoxazole, also known as cotrimoxazole, with Septrin and Bactrim being the common trade names.

Data have revealed agents that were previously unknown to be anti-Candida agents, which allows for the design of novel therapies against invasive candidiasis (14). Quinolones and other antibiotics can enhance the activity of azole and polyene agents; this synergistic action could have a clinically significant effect (15). Several drugs such as potassium iodide, amphotericin B, ketoconazole, itraconazole, and fluconazole as well as cotrimoxazole have been used successfully in the treatment of infections caused by *B. ranarum (16)*. For its antifungal use, it is acceptable but the application of cotrimoxazole requires *in vitro* assessment.

In view of the limited reports on the effect of “non-antifungal drugs”, further *in vitro* verification of these studies is necessary. The present study aimed to evaluate the *in vitro* antifungal effect of cotrimoxazole, artesunate, and chloramphenicol against *Basidiobolus* species.

## Methods

### Preparation of fungal strains

*Basidiobolus* species (n =16), which have been isolated from human gastrointestinal basidiobolomycosis (GIB) and lizards, were evaluated in this study. Details of the source of the strains are shown in Table 1.

**Table 1.** *Basidiobolus* species isolated from human gastrointestinal basidiobolomycosis and gecko lizards used in the antimicrobial assay.

| SN | LABORATORY<br>AND (DSM)CODE | IDENTITY | SOURCE | SENSITIVE (S) OR RESISTANT (R) AND INHIBITION ZONE (MM) |  |  |  |  |  |  |  |
| --- | --- | --- | --- | --- | --- | --- | --- | --- | --- | --- | --- |
|  |  |  |  | TMP/SMX | ART | CPL | AMB | FLC | ITZ | MTZ | VCZ |
| 1. | Doza | *Basidiobolus haptosporus-like\** | Human gastrointestinal basidiobolomycosis (GIB), Aseer region, Saudi Arabia (2013) | S (50) | S (75) | S (28) | R (0) | S (36) | S (65) | R (0) | S (45) |
| 2. | 9-4 | *B. haptosporus-like* | Human GIB, Aseer region, Saudi Arabia (2014) | S (48) | S (72) | S (30) | R (0) | S (32) | S (68) | R (0) | S (40) |
| 3. | F43-5 | *B. haptosporus-like* | Human GIB, Aseer region, Saudi Arabia (2016) | S (48) | S (70) | S (30) | R (0) | S (33) | S (60) | R (0) | S (43) |
| 4. | V81 (DSM06014) | *B. haptosporus-like* | Human GIB, Aseer region, Saudi Arabia (2017) | S (49) | S (73) | S (30) | R (0) | S (35) | S (62) | R (0) | S (44) |
| 5. | F15-1 | *B. haptosporus-like* | Human GIB, Aseer region, Saudi Arabia (2017) | S (50) | S (72) | S (30) | R (0) | S (34) | S (64) | R (0) | S (43) |
| 6. | F17-5 (DSM06015) | *Basidiobolus* sp. | Human GIB, Aseer region, Saudi Arabia (2017) | S (47) | S (71) | S (30) | R (0) | S (33) | S (65) | R (0) | S (42) |
| 7. | L1 (DSM107663) | *B. haptosporus-like* | Gecko lizard, Muhayil, Aseer region, Saudi Arabia (2017) | S (49) | S (74) | S (30) | R (0) | S (33) | S (62) | R (0) | S (44) |
| 8. | L2 (DSM107662) | *B. haptosporus-like* | Gecko lizard, Muhayil, Aseer region, Saudi Arabia (2017) | S (50) | S (73) | S (29) | R (0) | S (37) | S (64) | R (0) | S (45) |
| 9. | L3 (DSM 05995) | *Basidiobolus* sp. | Gecko lizard, Muhayil, Aseer region, Saudi Arabia (2017) | S (48) | S (74) | S (28) | R (0) | S (36) | S (65) | R (0) | S (42) |
| 10. | L4 | *Basidiobolus* sp. | Gecko lizard, Muhayil, Aseer region, Saudi Arabia (2017) | S (48) | S (73) | S (30) | R (0) | S (37) | S (63) | R (0) | S (44) |
| 11. | L6 | *Basidiobolus* sp. | Gecko lizard, Muhayil, Aseer region, Saudi Arabia (2017) | S (47) | S (72) | S (30) | R (0) | S (37) | S (62) | R (0) | S (43) |
| 12. | 891-10 | *Basidiobolus* sp. | Human GIB, Aseer region, Saudi Arabia (2018) | S (49) | S (73) | S (28) | R (0) | S (37) | S (63) | R (0) | S (44) |
| 13. | ACH230 | *Basidiobolus* sp. | Human GIB, Aseer region, Saudi Arabia (2020) | S (50) | S (74) | S (30) | R (0) | S (35) | S (65) | R (0) | S (43) |

**Abbreviations:** TMP/SMX, Co-trimoxazole (40/16 mg/ mL); ART, Artesunate (10 mg/ mL); CPL, Chloramphenicol (100 mg/ mL); AMB, Amphotericin B (50 mg/ mL); FLC, Fluconazole (2 mg/ mL); ITZ, Itraconazole (10 mg/ mL); MTZ, Metronidazole (5 mg/ mL); VCZ, Voriconazole (10 mg/ mL). S, sensitive; R, resistant.
DSM, Deutsche Sammlung von Mikroorganismen und Zellkulturen GmbH, Inhoffenstraße 7B, 38124 Braunschweig, Germany; \**Basidiobolus haptosporus*-like was identified in our previous study (6) and other strains were identified based on phenotypic properties and currently underway for full descriptions and designations.

The strains were obtained from stored stock at 4°C, then subcultured onto Sabouraud dextrose agar (SDA; Difco, Becton, Dickinson and Company, Sparks, Maryland) and incubated at 25°C for 10 days. Prior to antimicrobial testing, the viability and purity of each isolate were evaluated by microscopic examination. All procedures were performed within a class II biological safety cabinet in a biosafety level 3 laboratory.

### Preparation of antimicrobial agents

The following agents have been used in the study: cotrimoxazole (Bactrim, sulfamethoxazole and trimethoprim 150/ 15 mg/ mL; Roche, Basel, Switzerland), artesunate (10 mg/ mL, Yashica Pharmaceuticals Private Limited, Thane, Maharashtra, India); chloramphenicol (200 mg/ mL, Cloranfenicol, BioGer Laboratories in Caracas - Venezuela); amphotericin B (100 mg/mL, Sigma, Missouri, USA); fluconazole (2 mg/mL (Diflucan I.V. Roerig/Pfizer Inc., France); itraconazole (10 mg/mL, Sporanox I.V, Janssen Biotech N.V, Belgium); metronidazole (5 mg /mL, PSI Pharmaceutical Co., Jeddah, Saudi Arabia); voriconazole (10 mg/ mL, Vfend, Pfizer Inc.).

### Preparation of fungal suspensions (inocula)

Sterile normal saline was added to each agar slant, and the cultures were gently scraped with cotton swabs. The suspension containing conidia and hyphae was diluted 1:10 with RPMI 1640 medium (9). The suspension was transferred to a sterile tube and allowed to settle for 5 min. Then the transmittance of the upper homogeneous supernatant was read at 530 nm and adjusted to 95% transmittance (approximately 1 × 10^3^ to 5 × 10^3^ CFU/mL). The strains were tested against each antimicrobial alone to determine the minimum inhibitory concentrations (MICs). The procedures were repeated at least twice, and each fungal strain was tested in duplicate.

### Agar well diffusion method

All strains were screened for their susceptibility to the antifungal and non-antifungal drugs shown in Table 1 using the agar well diffusion method (17). The plates were inoculated with each of the test strains using a cotton swab impregnated with fungal suspension (approximately 1 × 10^3^ to 5 × 10^3^ CFU/mL). Subsequently, a 100 μL of each drug with the provided concentration (Table 1) was spotted into wells made in SDA plates. Plates were incubated at 30C for 3 to 5 days. The inhibition zone was next read, and a strain was recorded sensitive (S) or resistant (R) accordingly.

### Broth Microdilution method

The broth microdilution method (M38-A2) was done following the Clinical and Laboratory Standards Institute guidelines (18). The concentrations used to screen the inhibitory activities are shown in Table 1. A serial two-fold dilution of each antimicrobial agent was performed to determine the minimum inhibitory concentration (MIC).

MICs for AMB and azoles were defined as the lowest concentration of the drug at which there was no visible fungal growth (18). MIC of SMX/TMP is defined as the lowest drug concentration that caused 80% inhibition of visible fungal growth.

### Statistical analysis

To determine the mean difference between inhibitory measures (in triplicate) amongst the subject drugs, the nonparametric Wilcoxon Signed Rank Test was used for the analysis of antimicrobial combinations. *P*-value of < 0.05 was considered significant.

## Results

The inhibitory activity of artesunate, cotrimoxazole, chloramphenicol, and in comparison with standard antifungal antibiotics is shown in Table 1. The three tested drugs were found active (*in vitro*) against *Basidiobolus* species and were able to inhibit all Basidiobolus species strains’ growth.

### Determination of the minimum inhibitory concentrations

The minimum inhibitory concentrations (MIC) of artesunate, chloramphenicol, cotrimoxazole, and in comparison with voriconazole against *Basidiobolus* species are shown in Table 2. MIC value for artesunate was found to be 20 μg/ mL; for chloramphenicol, 3,130 20 μg/ mL; cotrimoxazole, 160 μg/ mL, and for voriconazole was 80 μg/ mL.

**Table 2.** Minimum inhibitory concentrations (MIC) of artesunate, chloramphenicol, cotrimoxazole, and in comparison with voriconazole against *Basidiobolus* species.

| Strain V81* | Artesunate (mg/ml) |  |  |  |  |  |  |  |  |  |
| --- | --- | --- | --- | --- | --- | --- | --- | --- | --- | --- |
|  | 5 | 2.5 | 1.25 | 0.63 | 0.31 | 0.16 | 0.08 | 0.04 | 0.02 | 0.01 |
|  | NG | NG | NG | NG | NG | NG | NG | NG | G | G |
| Strain V81 | Chloramphenicol (mg/ml) |  |  |  |  |  |  |  |  |  |
|  | 50 | 25 | 12.5 | 6.25 | 3.13 | 1.56 | 0.78 | 0.4 | 0.2 | 0.1 |
|  | NG | NG | NG | NG | G | G | G | G | G | G |
| Strain V81 | Cotrimoxazole (mg/ml) |  |  |  |  |  |  |  |  |  |
|  | 20/<br>8 | 10/<br>4 | 5/ 2 | 2.5/<br>1 | 1.25/<br>0.5 | 0.63/<br>0.25 | 0.31/<br>0.13 | 0.16/<br>0.7 | 0.08/<br>0.35 | 0.04/<br>0.18 |
|  | NG | NG | NG | NG | NG | NG | NG | G | G | G |
| Strain V81 | Voriconazole (mg/ml) |  |  |  |  |  |  |  |  |  |
|  | 5 | 2.5 | 1.25 | 0.63 | 0.31 | 0.16 | 0.08 | 0.04 | 0.02 | 0.01 |
|  | NG | NG | NG | NG | NG | NG | G | G | G | G |

\*Strain V81, *Basidiobolus haptosporus*-like (DSM06014); NG, no growth; G, growth.

### The combined effects of voriconazole with artesunate, cotrimoxazole, chloramphenicol against*Basidiobolus* species

The combined effect of voriconazole with artesunate, cotrimoxazole, chloramphenicol on *Basidiobolus* species are shown in Figs. 1 and 2.

**Fig. 1.**
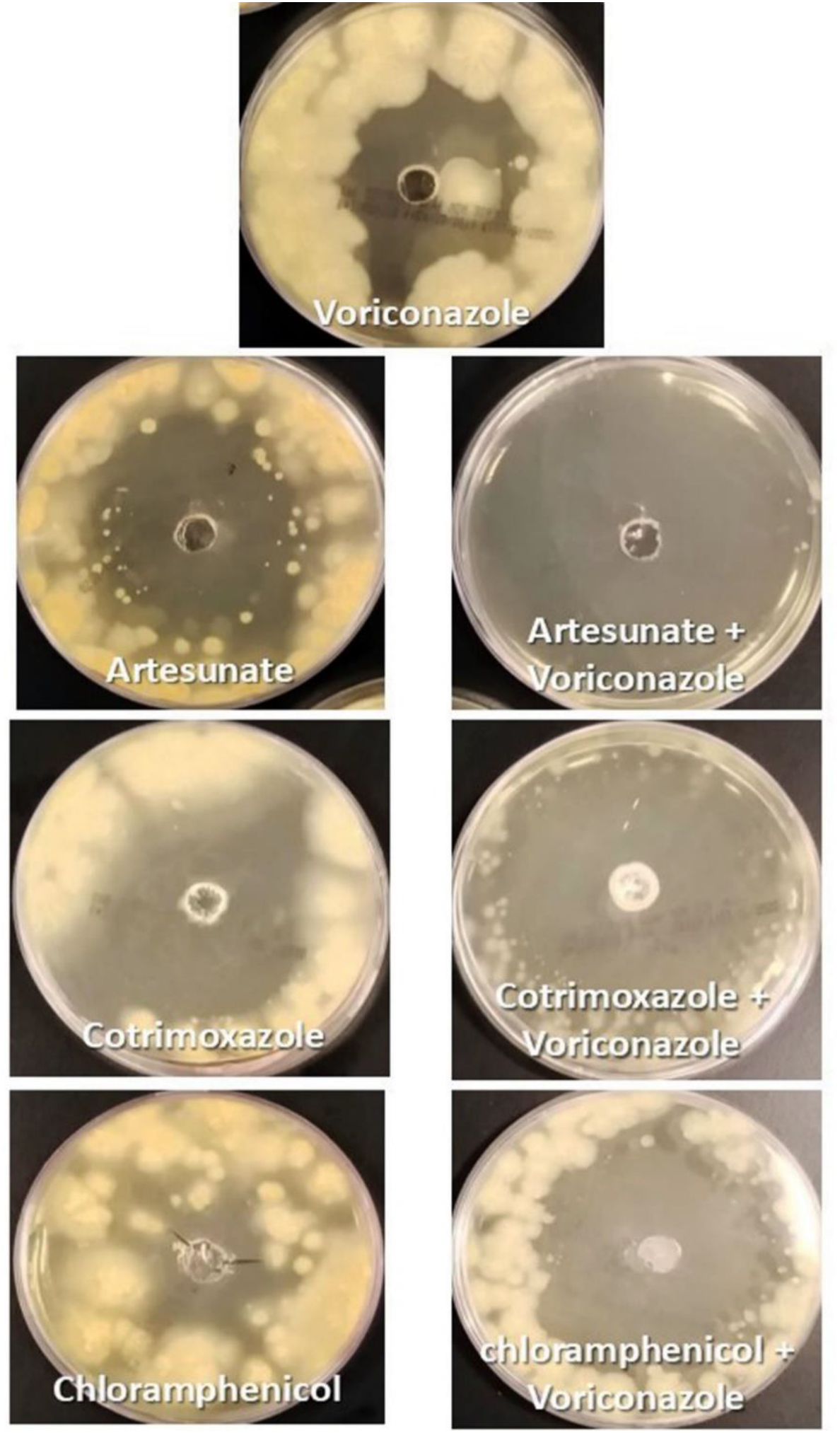
Zone of inhibition of the “non-antifungal" drugs on *Basidiobolus* sp. and in comparison with voriconazole.

**Fig. 2.**
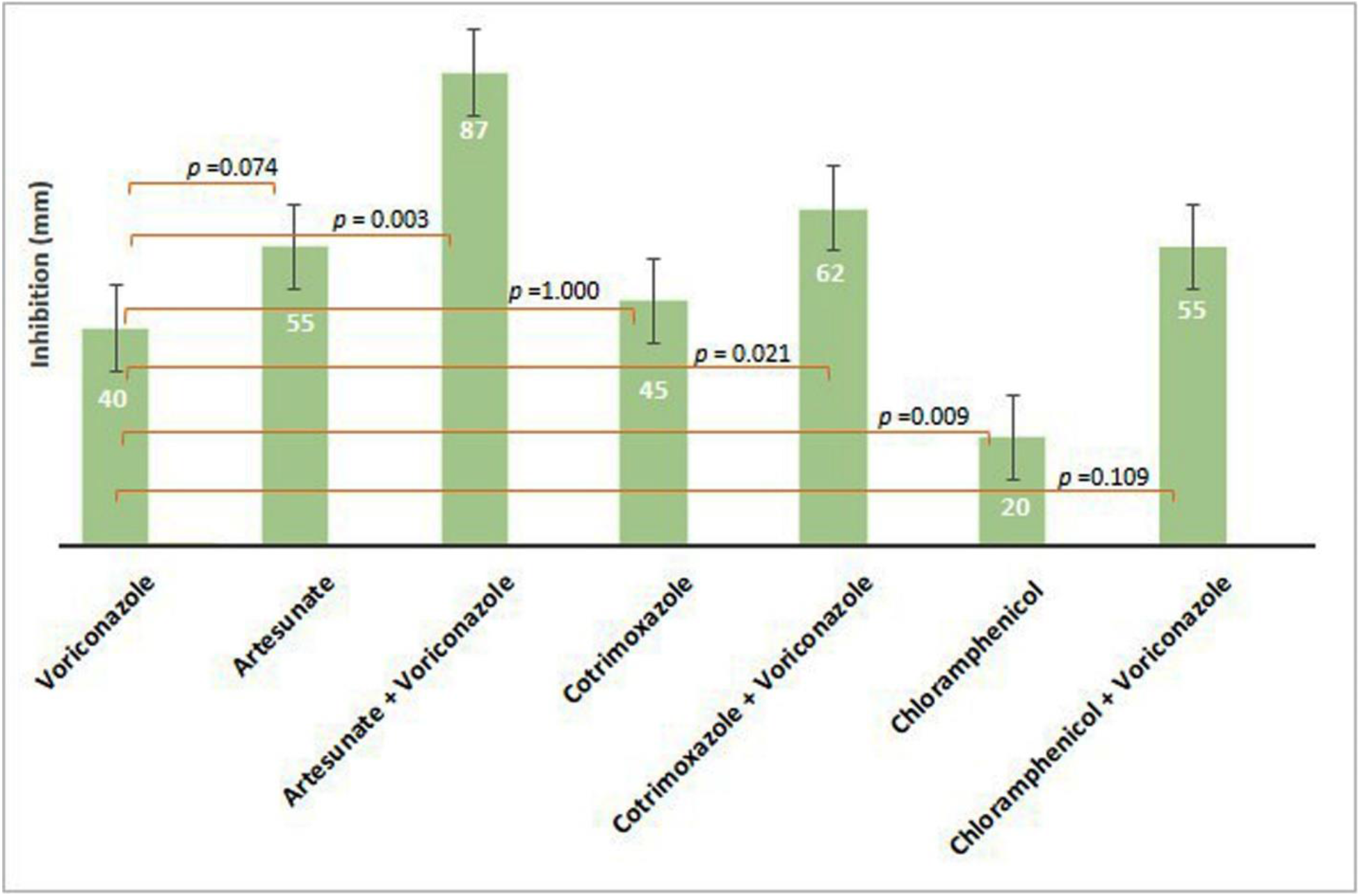
Comparison of the *in vitro* effect of voriconazole with “non-antifungal” drugs on *Basidiobolus* sp. (p <0.5 is considered significant).

The triplicate reading of the effect of voriconazole alone was 40-50 mm ±2.89. The effect of artesunate was 50-60 mm ± 2.89 compared to artesunate + voriconazole, 87-90 mm ± 1.45 (p = 0.003). The effect of cotrimoxazole was 40-50 mm ± 2.89 compared to cotrimoxazole + voriconazole, 60-64 mm ± 1.15 (p =0.021). The effect of chloramphenicol was 18-22 mm ± 1.15 compared to chloramphenicol + voriconazole, 53-57 mm ± 1.15 (p = 0.109).

Both artesunate and cotrimoxazole were as effective as voriconazole showing no significant difference between them, *p* = 0.074 and 1.0, respectively. Whereas, chloramphenicol was less active against *Basidiobolus* sp. (p= 0.009) (Fig. 2).

## Discussion

The results of our current study revealed the potential *in vitro* effect of artesunate, chloramphenicol, and cotrimoxazole as antifungal, in particular against *Basidiobolus* spp., the etiological agent of basidiobolomycosis. This is not completely new since few studies have suggested this idea and but most trials have been carried out on a clinical-based treatment course of therapy (10, 19). Nonetheless, *in vitro* evaluation of antifungal drugs and sulfamethoxazole-trimethoprim against clinical isolates of *Conidiobolus lamprauges* has been done (20). *Conidiobolus lamprauges* is a member of the order *Entomophthorales,* a species phylogentically related to *Basidiobolus ranarum*, *Basidiobolus haptosporus* or *Conidiobolus coronatus*, the later species are agents implicated in skin and abdominal basidiobolomycosis (16, 21).

Several conventional antifungal drugs, for example, potassium iodide, cotrimoxazole, amphotericin B, ketoconazole, and itraconazole, have been tried for the treatment of entomophthoromycosis due to *Conidiobolus coronatus* or basidiobolomycosis due to *Basidiobolus ranarum* or *Basidiobolus haptosporus*-like fungi with variable results (22). Comparably, information derived from a number of investigations indicated that non-antifungal compounds supplement the action of conventional antifungal agents either through the elimination of natural resistance or through exhibiting in some way activity antagonizing certain fungal species (14, 15, 19).

These results confirm the hypothesis that cotrimoxazole has an antifungal inhibitory, as reported previously (9). This substantiating that folic acid blockade may be a potential antifungal target for *Basidiobolus* species. This is the first report of the antifungal potential of cotrimoxazole drug against this fungal pathogen.

Potassium iodide and cotrimoxazole were found to be simple and effective for basidiobolomycosis treatment (23). While there is no consensus, African physicians prefer to use potassium iodide or trimethoprim-sulfamethoxazole in the treatment of tropical mycosis infections caused by either *Basidiobolus ranarum, Basidiobolus haptosporus* or *Conidiobolus coronatus* or others (24). Experience lead some physicians to adopt septrin (trimethoprim-sulfamethoxazole) as the drug of choice in the management of entomophthorosis due to *Conidiobolus coronatus* (25). Another study suggested using the combination of itraconazole and fluconazole as an additional option for the treatment of this mycosis acting better than sulfamethoxazole plus trimethoprim for 2 months (26). A case of rhinophycomycosis entomophthora was successfully treated with a combination of bacteria, potassium iodide, and steroid were reported (27). Clinical isolates of *Conidiobolus lamprauges* (entomophthoromycosis cases) showed synergistic interactions of 100% for the sulfamethoxazole-trimethoprim combination, 71% for the terbinafine-azole antifungal combination, and 29% for the terbinafine-micafungin combination. All other interactions were indifferent (20).

The mode of action of these three drugs is known against their conventional target organisms. It is, however, that the effect on fungi is still unknown. The inhibition of folic acid synthesis in *Coccidioides posadasii* has been speculated as a likely antifungal target (9). *Saccharomyces cerevisiae* in response to treatment with arsenic has revealed a complex response that influences signaling pathways, including protein kinases such as the mitogen-active protein kinase and target of the rapamycin complex 1 system (28). The modes of action of artesunate continue to be unclear and debatable (29).

On the other hand, chloramphenicol is a bacteriostatic that acts via inhibiting protein synthesis. It prevents protein chain elongation by inhibiting the peptidyl transferase activity of the bacterial ribosome (30). However, not known how it might affect the eukaryotic fungi.

Our data support the fact that artesunate is potential antifungal as well as its primary antimalarial activity (12). Data have suggested the enhancement of artesunate with miconazole in antagonizing *Candida’s* biofilm. These refer to the potential mixture treatment of *Candida albicans* biofilm-related infections. The results of our current study also validated the “non-antifungal antibacterial drugs as effective against *Basidiobolus* strains, which support their earlier antifungal effects (11, 13). The combined effect of artesunate + voriconazole and that of cotrimoxazole + voriconazole have significant synergic effects, p = 0.003 and p = 0.021, respectively. These are hopeful results for the treatment of GIB in humans, which need in vivo application and determining the clinical implications.

In conclusion, the results of the present study demonstrated a clear inhibitory antagonism of artesunate, cotrimoxazole, and a less significant effect in the case of chloramphenicol, against strains of *Basidiobolus*. These are encouraging results for the *in vivo* application on human or animal models. The *in vivo* application is necessary to be combined with standard antifungal drugs rather than a single therapy since our results have indicated a synergistic effect between cotrimoxazole and voriconazole, with a lesser effect between chloramphenicol and voriconazole, but markedly between artesunate and voriconazole. The study recommends the application of artesunate + voriconazole or cotrimoxazole + voriconazole in a clinical trial.

